# N100 as a Neural Marker of Atypical Early Auditory Encoding in Autism: Sensitivity to Pitch, Distance-Based Intensity, and Spatial Location

**DOI:** 10.1101/2025.08.08.669315

**Authors:** Sara Sharghilavan, Leila Mehdizadeh Fanid, Oana Geman, Hassan Shahrokhi, Hadi Seyedarabi

**Affiliations:** Department of Cognitive Neuroscience, Faculty of Education and Psychology, University of Tabriz,Tabriz, Iran; Data Science and AI, Computer Science and Engineering Department, Chalmers University of Technology; Data Science and AI, Computer Science and Engineering Department, University of Gothenburg, Gothenburg, Sweden; Data Science and AI, Computer Science and Engineering Department, Stefan cel Mare University of Suceava, Suceava, Romania; Autism and Related Neurodevelopmental Disorders Research Team, Tabriz University of Medical Sciences, Tabriz, Iran; Faculty of Electrical and Computer Engineering, University of Tabriz, Tabriz, Iran

**Keywords:** Autism, N100, Speech Processing, Pitch Processing, Spatial Processing

## Abstract

**Background:** Individuals with Autism Spectrum Disorder (ASD) show atypical auditory perception. The N100 event-related potential (ERP) reflects early auditory encoding, predictive coding, and sensory gain. Therefore, this study examined N100 responses to speech stimuli as a neural marker of auditory processing differences in ASD.

**Methods:** Event-related potentials (ERPs) were recorded using OpenBCI in 12 boys diagnosed with Level 1 ASD (requiring minimal support) and 15 typically developing (TD) peers. Participants passively listened to Romanian sentences systematically varied in pitch (normal, high, low), distance-based intensity (0.5, 1, 2 meters; 65, 59, 53 dB), and spatial presentation (binaural, left, right). N100 amplitudes and latencies were analyzed using Python and SPSS.

**Results:** ASD group indicated significantly reduced N100 amplitudes for normal-pitch stimuli (*p* = .030, η² = .175) and binaural presentation (*p* = .030, η² = .175). Marginal reductions were also observed for low pitch (*p* = .096, η² = .120), speech presented from a 0.5-meter distance (*p* = .058, η² = .147), and unilateral conditions (*p*s = .066–.077, η²s = .130–.142). No group differences emerged for N100 latency. These findings suggest attenuated early auditory responses in ASD to both typical and spatially complex speech cues.

**Conclusions:** Results support predictive coding models proposing reduced sensory precision in ASD. The consistent amplitude attenuation, including near-significant findings, points to subtle but pervasive impairments in early auditory encoding. The use of ecologically valid speech stimuli and portable EEG underscores the translational potential of N100 as a biomarker for early identification and intervention in autism.

## Introduction

Autism Spectrum Disorder (ASD) is a neurodevelopmental disorder that affects individuals across the entire lifespan (LeBlanc, Riley & Goldsmith, 2008; Tantam, 2012; Lai & Weiss, 2017; Tafolla, Singer & Lord, 2025). Two core diagnostic criteria for individuals with ASD include persistent deficits in social communication and interaction, and the presence of restricted, repetitive patterns of behavior, interests, or activities (American Psychiatric Association, 2013). Additionally, individuals with ASD have marked impairments in cognitive functions (Towgood, Meuwese, Gilbert, Turner & Burgess, 2009; Nicholl *et al*., 2014; Banker, Gu, Schiller & Foss-Feig, 2021), especially perception, processing, and attention (e.g., Liss, Saulnier, Fein & Kinsbourne, 2006; Marco, Hinkley, Hill & Nagarajan, 2011; Robertson& Baron-Cohen, 2017; Crasta, Salzinger, Lin, Gavin & Davies, 2020; Hadad & Yashar, 2022).

One of the core domains of impairment in ASD individuals is auditory processing (O’Connor, 2012; Rotschafer, 2021), which frequently involves atypical responses to auditory stimuli, ranging from hypersensitivity (Gomes, Pedroso, & Wagner, 2008; Lucker, 2013; Ida Eto, Hara, Ohkawara & Narita, 2017; de Albuquerque Britto, de Santana, Dias & da Silva Júnior, 2024) to hyposensitivity (Tan, Xi, Jiang, Shi, Wang & Wang, 2012; Neklyudova, Smirnov, Rebreikina, Martynova & Sysoeva, 2022). Given that auditory processing forms the foundation of speech processing (Nelken, 2008; Price & Moncrieff, 2021), and considering that Speech is a complex cognitive function that integrates both acoustic and linguistic elements (Sharghilavan, Geman, Toderian, 2024), a thorough investigation of early-stage auditory processing is essential to elucidate the mechanisms underlying speech perception. Therefore, the present study investigates differences in the auditory processing of speech-based stimuli manipulated in pitch, spatial direction, and temporal speed between individuals with high-functioning autism and typically developing individuals.

Auditory processing differences are increasingly recognized as foundational in ASD, yet their underlying neural mechanisms remain incompletely understood. Neurophysiological tools such as continuous EEG monitoring (Mihai et al., 2025) and event-related potentials (ERPs) provide critical insights into both early and higher-order processing (Jeste & Nelson, 2009; Cotter et al., 2023). Among these, the N100 (or N1) component—peaking approximately 80–120 ms post-stimulus in fronto-central regions—is a robust marker of early auditory encoding (Rosburg, Boutros, & Ford, 2008). N100 is reliably elicited even in passive listening (Handy, 2005; Kappenman & Luck, 2016; Ren et al., 2021), and its amplitude varies with stimulus type and intensity, reaching up to −5 µV for highly intense sounds (Näätänen & Picton, 1987; Sharma & Dorman, 2000).

Despite its utility, the influence of specific acoustic features such as pitch, intensity, and spatial direction on the N100 remains underexplored in ASD. The broader literature reports inconsistent findings: while some studies suggest superior pitch discrimination in ASD (Bonnel et al., 2003; Heaton et al., 2008; Chowdhury et al., 2017; Chen et al., 2022), others report pitch perception deficits (Bhatara et al., 2013; Ong et al., 2024). Similarly, sensitivity to sound intensity shows conflicting patterns, with both hyper- and hypo-responsiveness reported (Khalfa et al., 2004; Jones et al., 2009; Hsieh et al., 2022; Bruneau et al., 2003), and findings in spatial attention remain equivocal (Teder-Sälejärvi et al., 2005; Soskey et al., 2017). Critically, no prior studies have systematically examined how children with ASD neurally encode these core acoustic features of speech—pitch, intensity, and spatial location—as reflected in the N100.

To address this gap, the present study investigates early auditory cortical responses to ecologically valid speech stimuli systematically varied across pitch, intensity (via source distance), and spatial direction. Using a portable, child-friendly EEG system, we compare N100 amplitudes and latencies between children with ASD and their typically developing peers. By characterizing neural responses to these fundamental auditory dimensions, the study aims to clarify early-stage encoding mechanisms in autism and inform predictive models of atypical sensory processing (Nelken, 2008; Okada et al., 2010; Price & Moncrieff, 2021).

## Materials and Methods

### Participants

We recruited 12 boys with high-functioning autism [mean age = 9.7 years, range = 7–12 years] and 15 typically developing boys [mean age = 9.3 years, range = 7–12 years]. Participants in the two groups were individually matched for chronological age, intelligence, and handedness. All participants were native Romanian speakers and demonstrated normal hearing, as confirmed by medical records indicating hearing thresholds of 25 dB or better across frequencies ranging from 250 to 8000 Hz. The diagnosis of ASD was based on clinical assessments and formal diagnosis using the Autism Diagnostic Interview-Revised (ADI-R) (Lord, Rutter, & Le Couteur, 1994). No participants had a history of neurological or psychiatric comorbidities. Before participation, written informed consent was obtained from the parents of all participants, following the Declaration of Helsinki (World Medical Association, 2003). The study protocol was approved by the Research Ethics Committees of tefan cel Mare University and the University of Tabriz (IR.TABRIZU.REC.1403.172).

### Stimuli

This experiment formed a subset of a broader research project. A corpus of brief Romanian sentences simply describing objects was employed. Each auditory stimulus comprised short three-word sentences, recorded at a natural speech rate (∼3.5 syllables per second) within the contralto vocal range (F3) (approximately 174 Hz) under noise-free studio conditions. To ensure consistency across stimuli, all recordings were normalized by applying a uniform fundamental frequency (F0), thereby maintaining a stable and homogeneous acoustic profile. This normalization was intended to reduce inter-speaker and physiological variability that could otherwise confound neural and auditory responses. Following this, specific acoustic parameters, most notably loudness as a proxy for perceived source distance, were systematically manipulated to support the experimental objectives. Subsequently, the acoustic features were manipulated to align with the study’s objectives, including spatial distance, spatial direction, and pitch.

Pitch manipulation involved adjusting the fundamental frequency by one whole tone (two semitones), high or low pitch. This magnitude of change was selected to be perceptible while preserving the naturalness of speech, thus maintaining validity for cognitive processing analyses.

**Figure 1.**
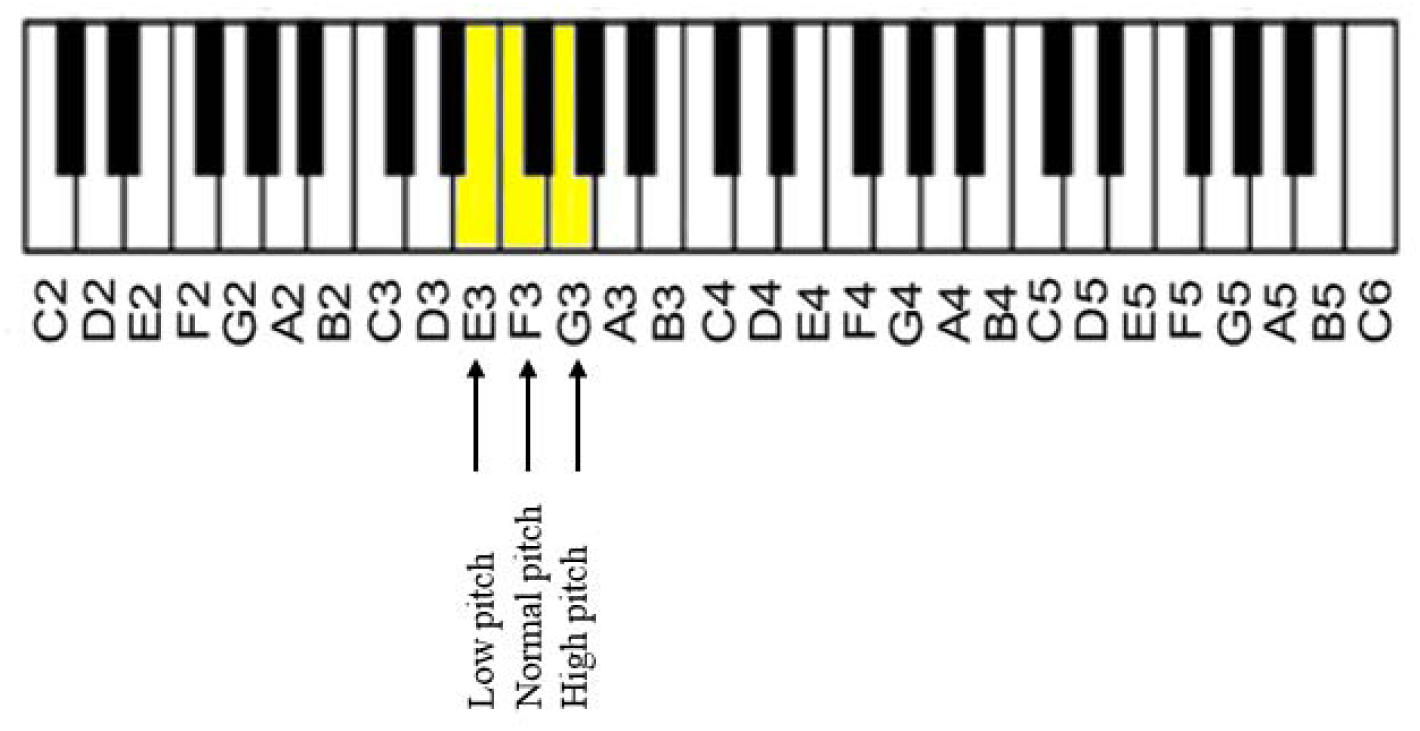
Pitch presentation conditions (normal, relatively high, and relatively low pitch) of auditory stimuli

For manipulation of distance, we considered a sound level of 65 dB SPL at a reference distance of half a meter (Pearsons, Bennett & Fidell, 1977). Using the free-field sound attenuation formula derived from the inverse square law (Martinis & Ozimec, 2011):

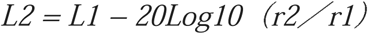

To achieve a perceptually realistic simulation of auditory distance, a multifaceted approach integrating several acoustic cues was implemented, extending beyond mere intensity adjustments. Specifically, according to the formula mentioned above, sound pressure levels of 65, 59, and 53 dB SPL were systematically assigned to represent source distances of 0.5 m, 1 m, and 2 m, respectively, in accordance with free-field attenuation principles. In addition, to enhance ecological validity, spectral cues were incorporated by attenuating high-frequency components in stimuli corresponding to greater distances, thereby emulating natural acoustic filtering during sound propagation. Reverberation characteristics appropriate for each virtual distance were simulated using convolution with empirically derived room impulse responses, enabling systematic manipulation of the direct-to-reverberant energy ratio (DRR). Furthermore, binaural rendering via headphones preserved spatial fidelity by delivering precise interaural cues. Finally, the auditory output was calibrated using Room EQ Wizard (REW) software in combination with Apple AirPods Pro. Standardized test signals were employed, and output levels were fine-tuned to ensure accurate reproduction of the target SPL values. Also, to manipulate direction, we played the sentences monaurally, through either the right or left ear, as opposed to the standard sentences that were presented binaurally. Auditory stimuli were binaurally rendered and simultaneously presented through headphones to both ears, simulating a sound source at a 90-degree horizontal azimuth.

**Figure 2.**
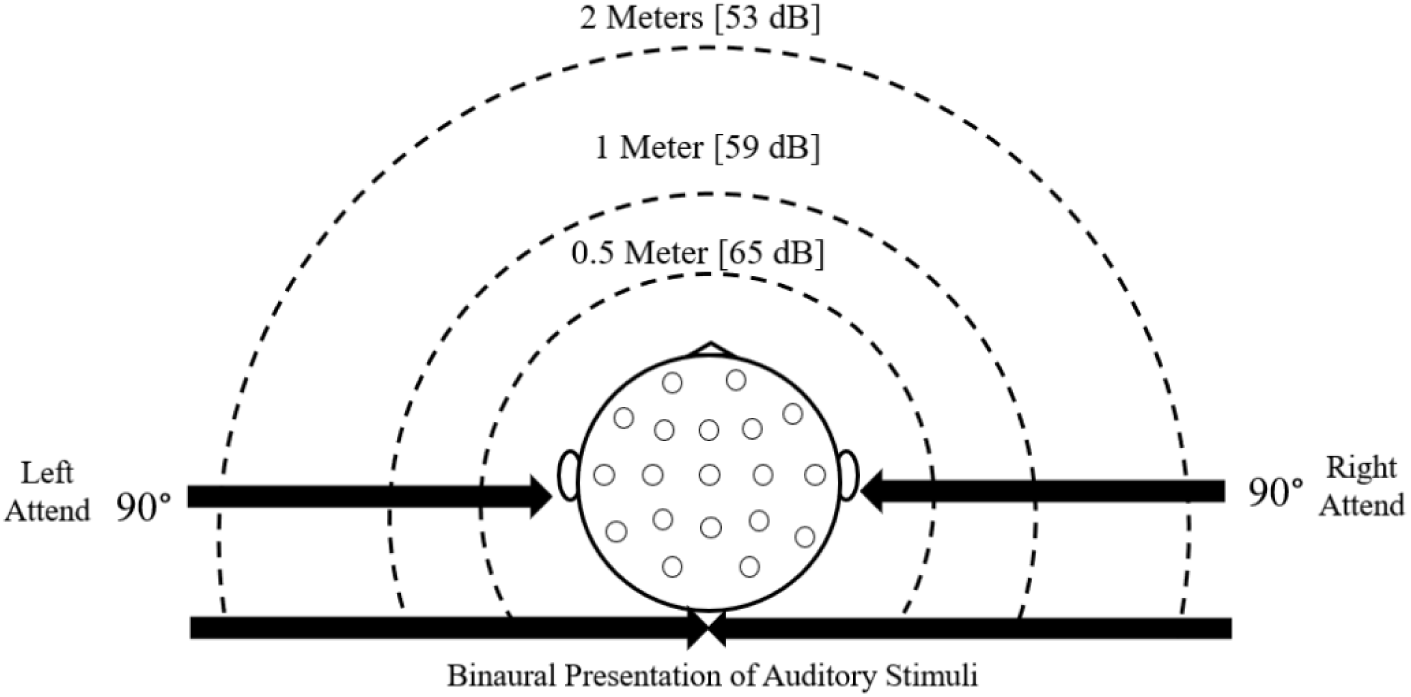
Spatial presentation conditions (distance and direction) of auditory stimuli Apparatus

Electroencephalography (EEG) is a fundamental method for monitoring brain activity (Mihai, Geman, Toderean, Miron & SharghiLavan, 2025). Techniques such as event-related potentials (ERPs), which extract specific brain responses, have significantly advanced our understanding of brain function. ERPs are changes in EEG signals triggered by sensory stimuli exposure (Barry, De Blasio, Rushby, MacDonald, Fogarty, & Cave, 2022). In the present study, we employed the Ultracortex Mark IV EEG headset, developed by OpenBCI, as the brain-computer interface (BCI). This device features 16 dry electrodes arranged according to the international 10-20 system, ensuring comprehensive coverage of key cortical areas. The headset interfaces with the Cyton board, an advanced biosensor platform capable of capturing EEG, EMG, and ECG signals. Data are sampled at 250 Hz and transmitted wirelessly to a computer via an RFduino Bluetooth module connected through a USB dongle. This wireless setup minimizes movement constraints and is particularly advantageous for experiments involving children or conducted in low-stimulation environments (OpenBCI, 2024).

### Experimental Procedure

Following initial screening, eligible participants underwent individual EEG recording sessions in a quiet, controlled environment. Seated comfortably with eyes closed to minimize artifacts, participants were fitted with an OpenBCI EEG headset using the 10–20 electrode placement system. Impedance was maintained below 10 kΩ, and brain activity was recorded via the OpenBCI Cyton Board.

Since this study is part of a huge project, the stimuli were presented in an oddball task which presented binaurally, with standard stimuli (e.g., normal pitch, 65 dB SPL without any distance cue, binaural presentation) occurring on 75% of trials, and deviant stimuli (e.g., ±2 semitones in pitch, lower SPLs, or lateralized direction) on 25%. To ensure that participants were attentively listening to the auditory stimuli, a brief comprehension check was administered following the experiment, during which participants were asked questions related to the content of the presented sentences. All stimuli were presented at the sentence level, but ERP analysis focused on the initial syllables, with triggers aligned at sentence onset to ensure temporal precision.

A custom Python interface controlled stimulus presentation and synchronized ERP event markers with the EEG recording (Sharghilavan, Geman, & Toderean, 2024), enabling accurate analysis of neural responses to speech stimuli.

**Figure 3.**
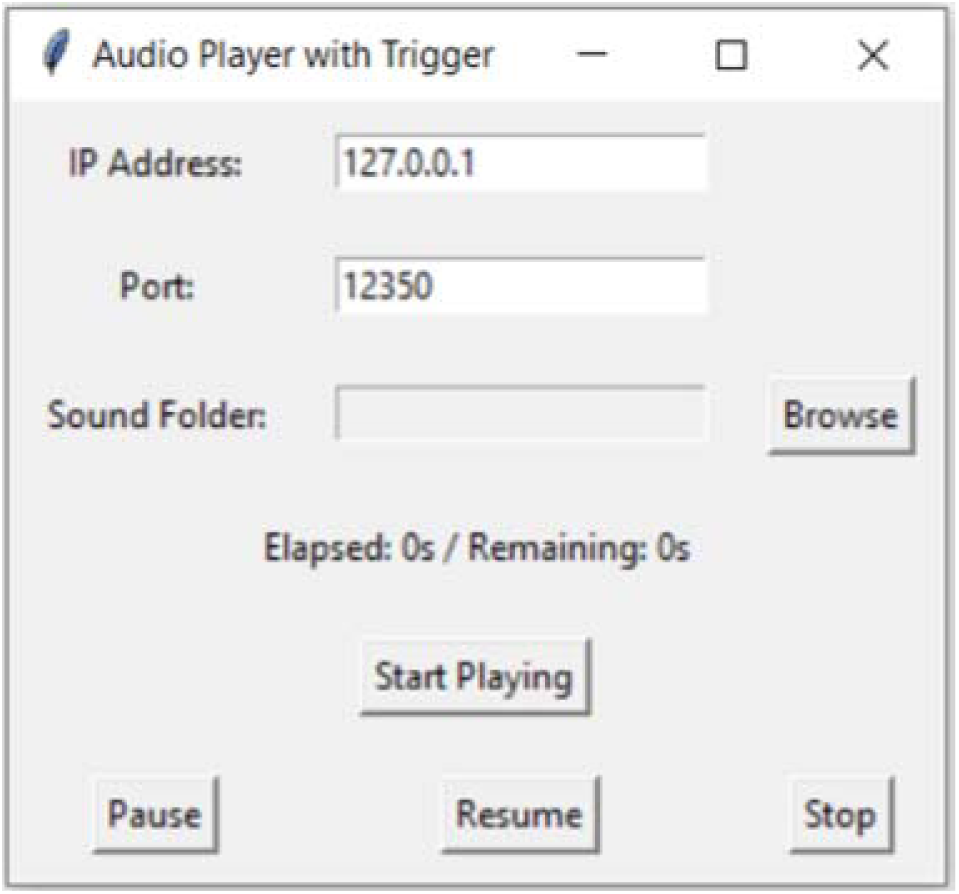
Custom Python interface for this project

Following data acquisition, EEG signals containing ERP markers were processed using Python, employing the Pandas, Numpy, and Matplotlib libraries for data handling, statistical analysis, and visualization.

### Data analysis

#### Python

EEG signals were acquired in BDF format, incorporating event-related potential (ERP) markers for time-locked analysis. Data preprocessing and analysis were performed in Python using libraries, including NumPy, Pandas, and Matplotlib. To enhance signal fidelity and minimize noise interference, a fourth-order Butterworth bandpass filter with a passband of 0.1–40 Hz was applied, effectively attenuating low-frequency drifts and high-frequency artifacts. Artifact removal was further refined via Independent Component Analysis (ICA), allowing for the identification and exclusion of components associated with ocular and myogenic artifacts while preserving neurophysiologically relevant sources. Baseline correction was implemented by subtracting the mean voltage within a 200-ms pre-stimulus window from each epoch to compensate for slow potential shifts unrelated to stimulus onset.

Stimulus-locked epochs were segmented from −300 ms to +1000 ms relative to stimulus onset. Trials with missing or corrupted segments were excluded from further analysis. For each valid epoch, ERP features were extracted by computing amplitude (peak-to-peak mean voltage) and latency (time point of maximal or minimal deflection within the defined component window). These parameters facilitate subsequent multivariate statistical analyses across experimental conditions and cognitive domains, as elaborated in the Results section.

#### SPSS

Statistical analyses were performed using IBM SPSS Statistics version 27.0.1 (IBM Corp., Armonk, NY, USA). A multivariate analysis of variance (MANOVA) was conducted to examine the effects of group (ASD vs. TD) on the dependent variables, which included both peak amplitude and latency of the N100 in all conditions. Before conducting MANOVA, assumptions of multivariate normality, homogeneity of variance-covariance matrices (Box’s M test), and absence of multicollinearity were assessed.

## Results

Before conducting the MANOVA, all necessary statistical assumptions were rigorously examined. Univariate normality was verified using the Kolmogorov–Smirnov test, with all p-values exceeding the .05 threshold. Homogeneity of variance–covariance matrices was confirmed by a non-significant Box’s M test result. Linearity and absence of multicollinearity were assessed through visual inspection of scatterplots and correlation analyses, both indicating acceptable levels. Outlier detection based on standardized residuals and Mahalanobis distance revealed no influential cases. Given that all assumptions were satisfactorily fulfilled, the dataset was deemed suitable for MANOVA. The subsequent section outlines the results concerning group differences in ERP amplitude and latency.

### N100 Responses to pitch

**Figure 4.**
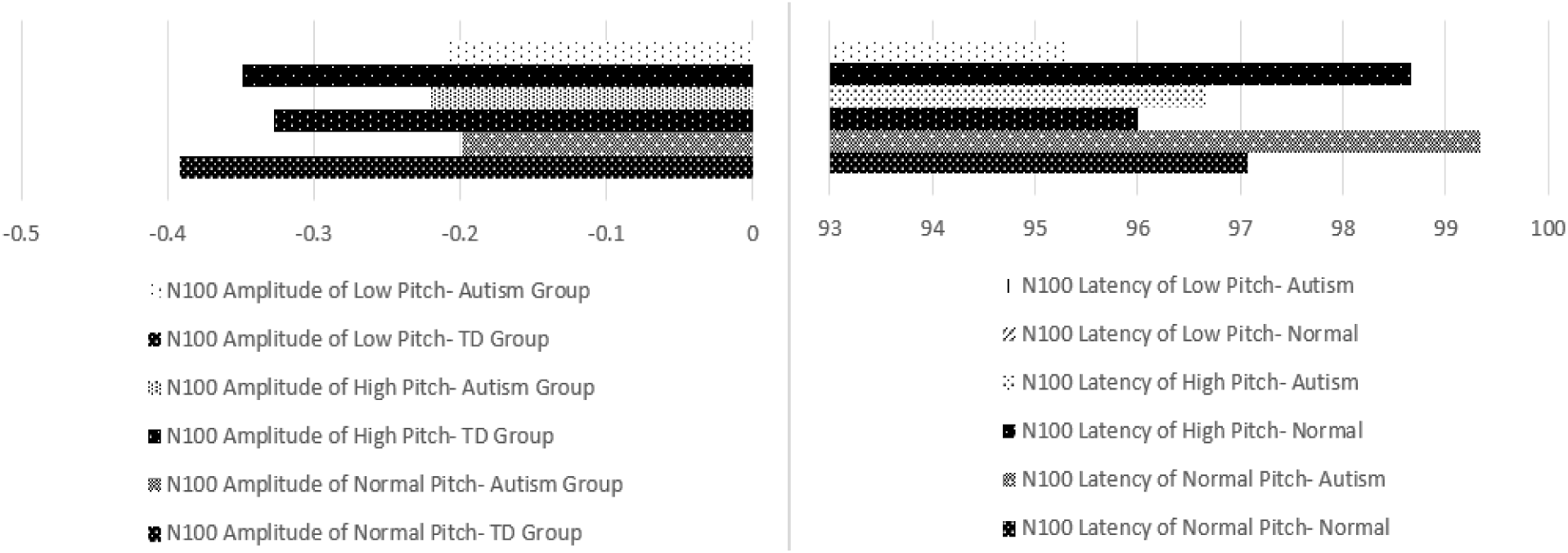
Comparison of N100 amplitude and latency between children with ASD and TD peers across three pitch deviation conditions (high, low, and normal).

Descriptive statistics revealed that the TD group exhibited consistently more negative N100 amplitudes across pitch conditions compared to the ASD group (normal: −0.39 µV vs. −0.20 µV; high: −0.33 µV vs. −0.22 µV; low: −0.35 µV vs. −0.21 µV), indicating reduced auditory cortical responses in ASD. Latency differences were small and variable: ASD latencies were slightly longer for normal pitch (99.33 ms vs. 97.07 ms), similar for high pitch (96.67 ms vs. 96.00 ms), and shorter for low pitch (95.33 ms vs. 98.67 ms).

**Table 1.** Summary of multivariate test results for the main effect of group on N100 amplitude and latency, considering all pitch deviation conditions combined.

|  |  |  |  |  |  |  | Partial Eta |
| --- | --- | --- | --- | --- | --- | --- | --- |
|  | Effect | Value | F | Hypothesis df | Error df | Sig. | Squared |
| Group | Pillai's Trace | .309 | 1.493 | 6.000 | 20.000 | .231 | .309 |
|  | Wilks' Lambda | .691 | 1.493 | 6.000 | 20.000 | .231 | .309 |
|  | Hotelling's Trace | .448 | 1.493 | 6.000 | 20.000 | .231 | .309 |
|  | Roy's Largest Root | .448 | 1.493 | 6.000 | 20.000 | .231 | .309 |

A one-way MANOVA was conducted to examine group differences (TD vs. ASD) across six N100 measures (amplitudes and latencies in normal, high, and low pitch conditions). The multivariate effect of group was not statistically significant, Wilks’ Lambda = .691, F(6, 20) = 1.493, p = .231, partial η² = .309, indicating no strong overall group effect. Although the p-value did not reach conventional significance, the partial eta-squared value suggests a moderate effect size, implying that group membership may account for a substantial portion of variance across measures. To further explore potential group differences, a between-subjects analysis was conducted. Results for N100 amplitude and latency across stimulus types are presented in Table 2.

**Table 2.**
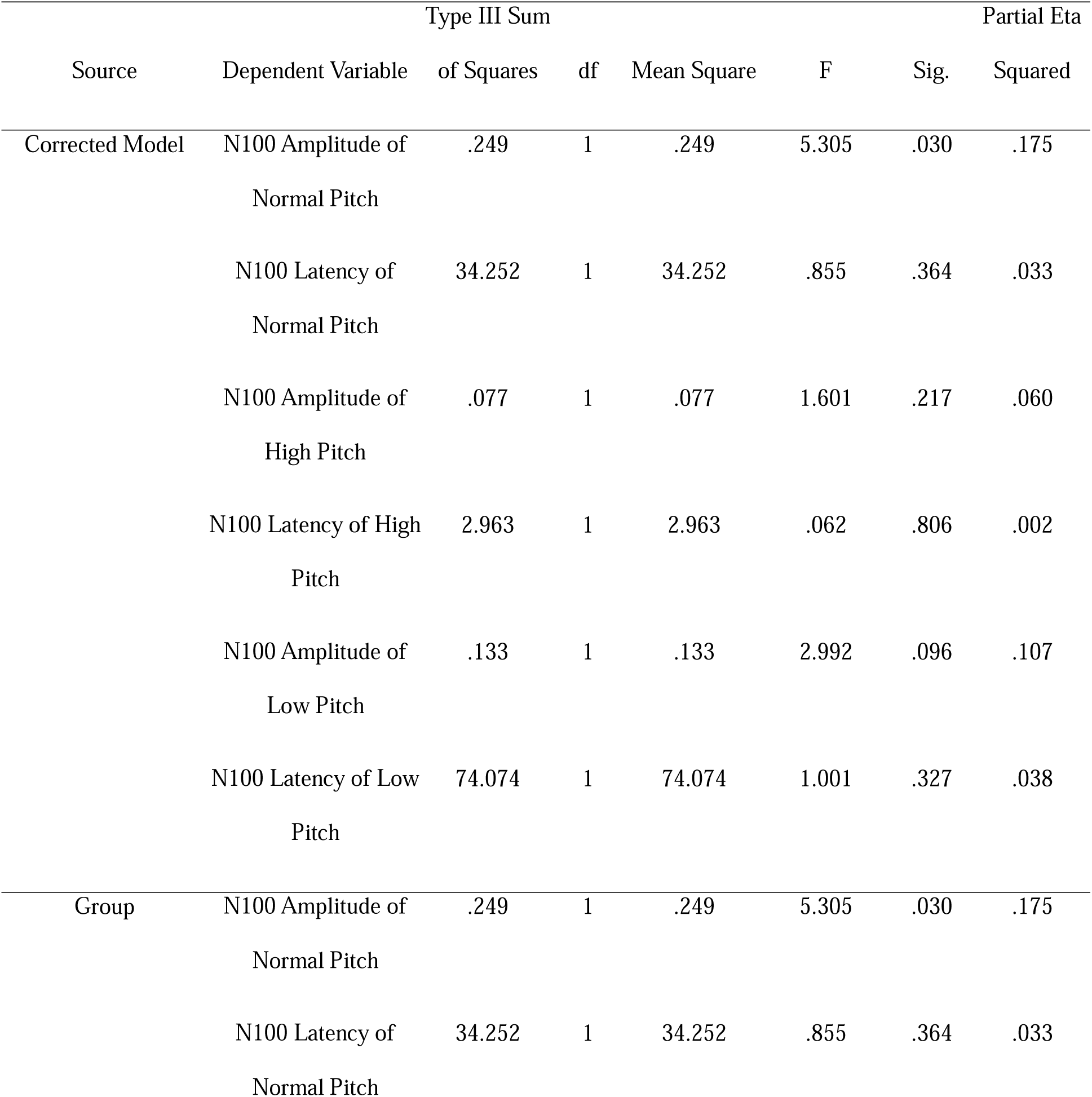
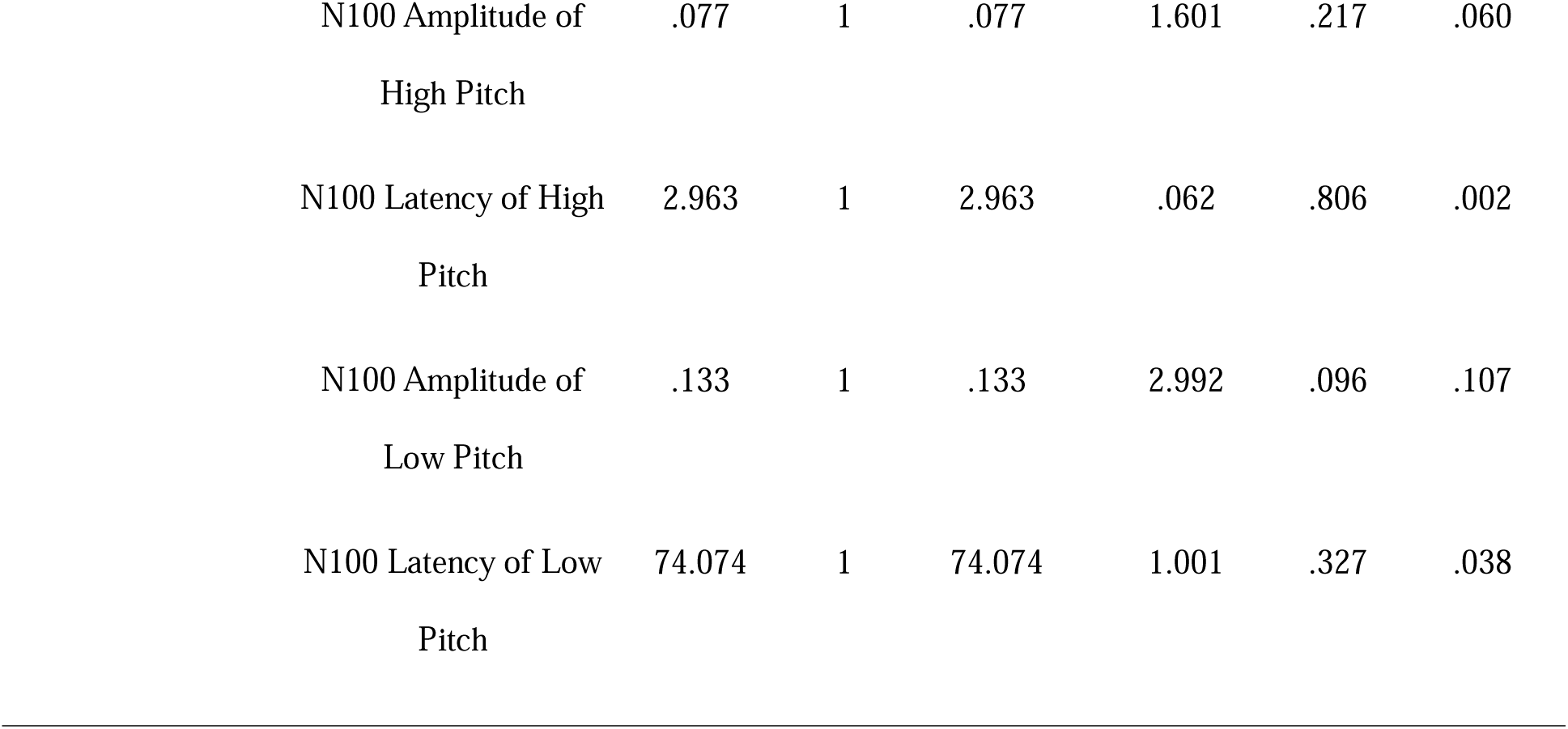
Tests of between-subjects effects on N100 amplitude and latency across pitch deviation conditions (normal, high, and low) comparing children with ASD and TD peers.

Between-subjects ANOVAs were conducted to examine group differences (ASD vs. Typically Developing) in the amplitude and latency of the N100 component across three pitch conditions (normal, high, and low). A significant group effect was observed for the amplitude of N100 in the normal pitch condition, F(1, 25) = 5.305, *p* = .030, partial η² = .175, indicating a moderate effect of group. In addition, the amplitude of N100 in the low pitch condition showed a trend toward significance, F(1, 25) = 2.992, *p* = .096, partial η² = .107, suggesting a possible group-related difference in auditory processing. No other comparisons reached statistical or trend-level significance. These findings suggest that group differences were most pronounced in response to normal-pitch stimuli and may also emerge in low-pitch conditions, particularly in amplitude measures.

### N100 Responses to Sound Intensity (Sound Distance)

**Figure 5.**
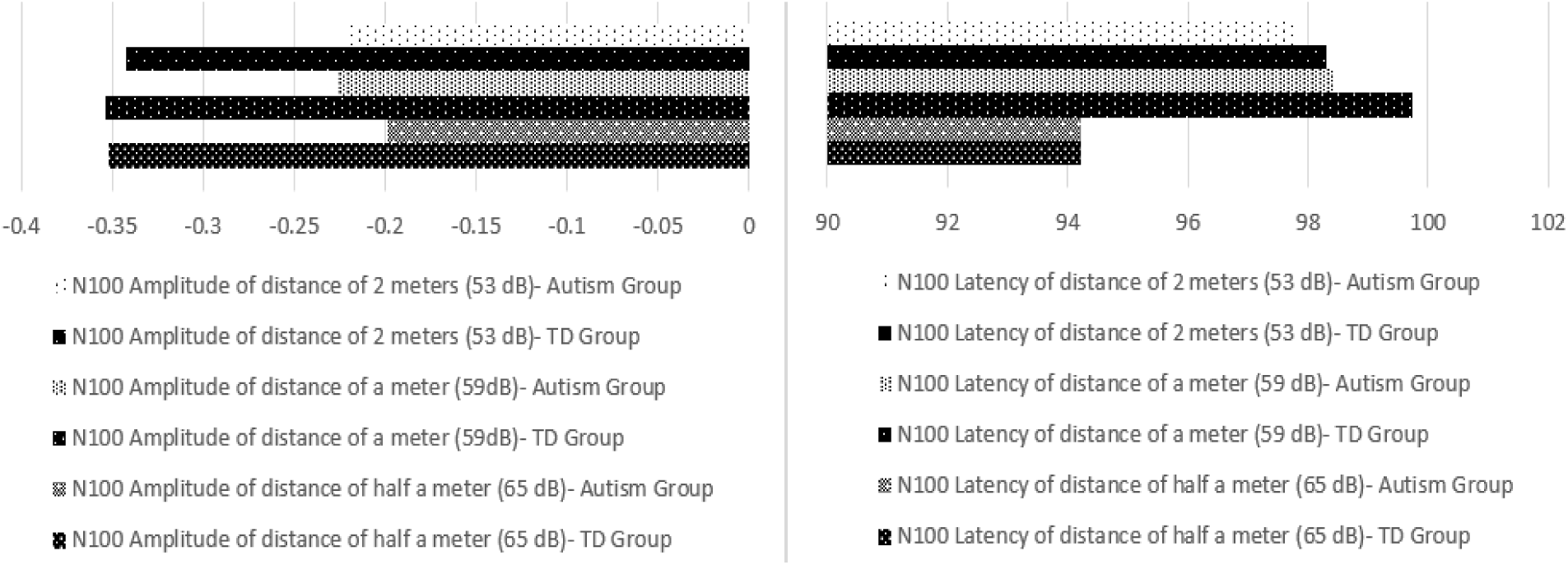
Comparison of N100 amplitude and latency between children with ASD and TD peers across three auditory distance conditions (2 meters, 1 meter, and 0.5 meter), reflecting varying levels of sound intensity.

Across all intensity levels (65, 59, and 53 dB; 0.5, 1, and 2 m), the ASD group showed consistently reduced N100 amplitudes compared to TD peers (e.g., at 65 dB: −0.35 µV TD vs. −0.20 µV ASD). In contrast, latencies were comparable between groups, with minimal variation across conditions. These findings suggest attenuated early auditory responses in ASD, primarily evident in amplitude rather than latency.

**Table 3.**
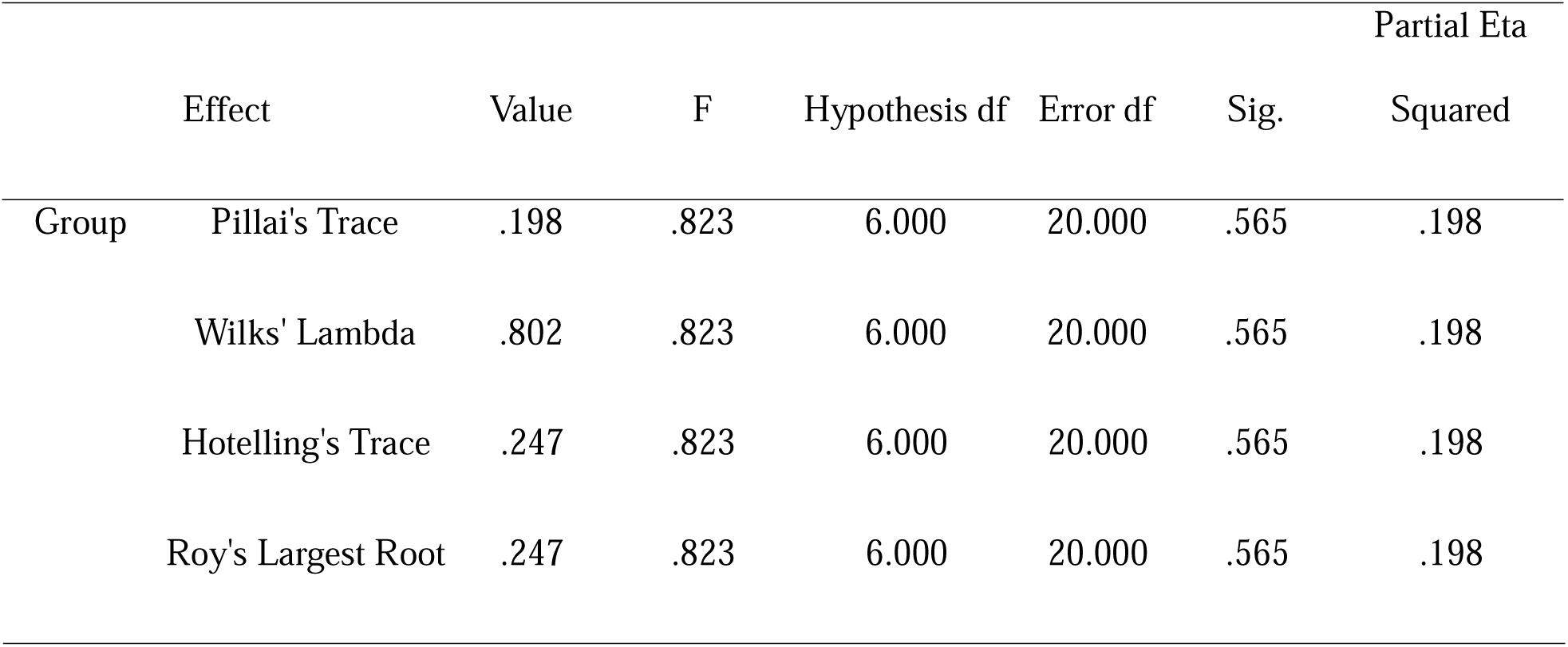
Summary of multivariate test results for the main effect of group on N100 amplitude and latency, considering all sound distance (sound intensity) deviation conditions.

|  |  |  |  |  |  |  | Partial Eta |
| --- | --- | --- | --- | --- | --- | --- | --- |
|  | Effect | Value | F | Hypothesis df | Error df | Sig. | Squared |
| Group | Pillai's Trace | .198 | .823 | 6.000 | 20.000 | .565 | .198 |
|  | Wilks' Lambda | .802 | .823 | 6.000 | 20.000 | .565 | .198 |

|  |  |  |  |  |  |  |
| --- | --- | --- | --- | --- | --- | --- |
| Hotelling's Trace | .247 | .823 | 6.000 | 20.000 | .565 | .198 |
| Roy's Largest Root | .247 | .823 | 6.000 | 20.000 | .565 | .198 |

Multivariate analysis of variance (MANOVA) was conducted to examine the effect of group differences across the dependent variables. The multivariate test statistics, including Pillai’s Trace, Wilks’ Lambda, Hotelling’s Trace, and Roy’s Largest Root, all yielded non-significant results, F(6, 20) = 0.823, p = .565, with a partial eta squared of .198. These findings indicate that there were no statistically significant multivariate effects of group on the combined dependent variables. Although the effect size, as indicated by partial eta squared, suggests a small-to-moderate proportion of explained variance, the lack of statistical significance implies that any group differences are not robust within the current sample. However, to further explore potential group differences, a between-subjects analysis was conducted. Results for N100 amplitude and latency across stimulus types are presented in Table 4.

**Table 4.**
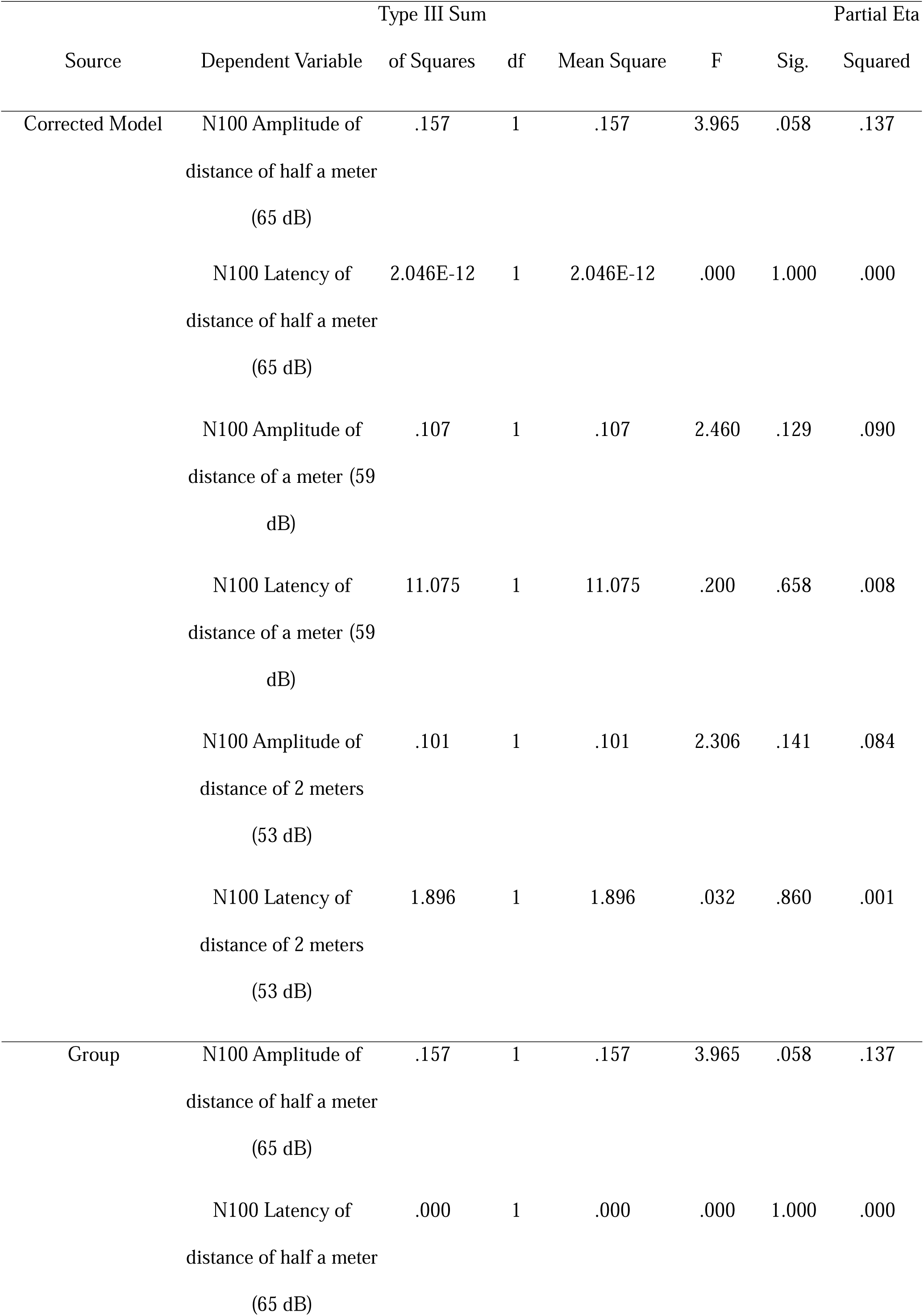
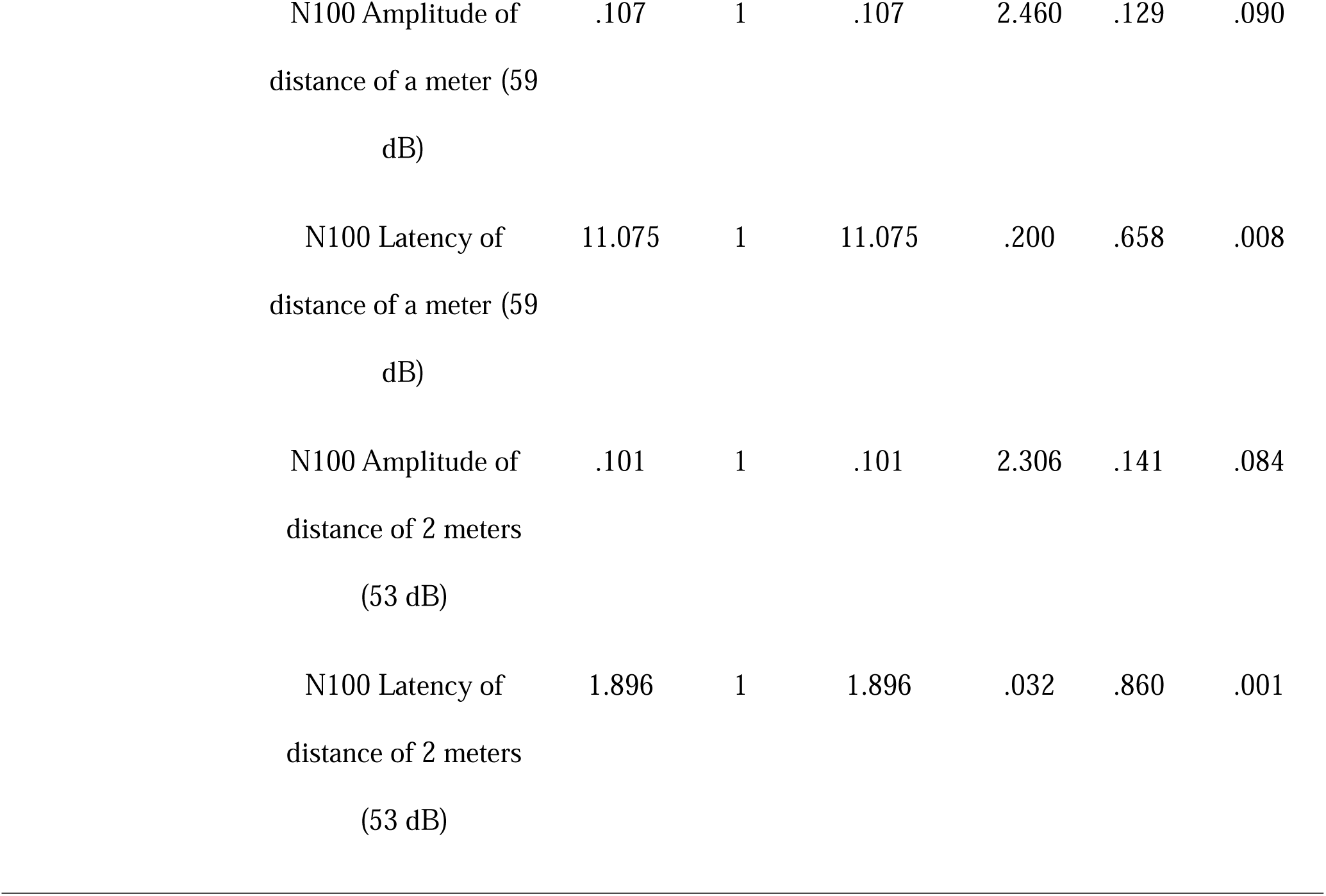
Tests of between-subjects effects on N100 amplitude and latency across sound distance (intensity) conditions comparing children with ASD and TD peers.

| Source | Dependent Variable | Type III Sum |  |  | F | Sig. | Partial Eta Squared |
| --- | --- | --- | --- | --- | --- | --- | --- |
|  |  | of Squares | df | Mean Square |  |  |  |
| Corrected Model | N100 Amplitude of<br>distance of half a meter<br>(65 dB) | .157 | 1 | .157 | 3.965 | .058 | .137 |

|  |  |  |  |  |  |  |
| --- | --- | --- | --- | --- | --- | --- |
| N100 Latency of | 2.046E-12 | 1 | 2.046E-12 | .000 | 1.000 | .000 |
| distance of half a meter |  |  |  |  |  |  |
| (65 dB) |  |  |  |  |  |  |
| N100 Amplitude of | .107 | 1 | .107 | 2.460 | .129 | .090 |
| distance of a meter (59 |  |  |  |  |  |  |
| dB) |  |  |  |  |  |  |
| N100 Latency of | 11.075 | 1 | 11.075 | .200 | .658 | .008 |
| distance of a meter (59 |  |  |  |  |  |  |
| dB) |  |  |  |  |  |  |
| N100 Amplitude of | .101 | 1 | .101 | 2.306 | .141 | .084 |
| distance of 2 meters |  |  |  |  |  |  |
| (53 dB) |  |  |  |  |  |  |
| N100 Latency of | 1.896 | 1 | 1.896 | .032 | .860 | .001 |
| distance of 2 meters |  |  |  |  |  |  |
| (53 dB) |  |  |  |  |  |  |
| Group | N100 Amplitude of | .157 | 1 | .157 | 3.965 | .058 | .137 |
|  | distance of half a meter |  |  |  |  |  |  |
|  | (65 dB) |  |  |  |  |  |  |

|  |  |  |  |  |  |  |
| --- | --- | --- | --- | --- | --- | --- |
| N100 Latency of | .000 | 1 | .000 | .000 | 1.000 | .000 |
| distance of half a meter |  |  |  |  |  |  |
| (65 dB) |  |  |  |  |  |  |
| N100 Amplitude of<br>distance of a meter (59<br>dB) | .107 | 1 | .107 | 2.460 | .129 | .090 |
| N100 Latency of<br>distance of a meter (59<br>dB) | 11.075 | 1 | 11.075 | .200 | .658 | .008 |
| N100 Amplitude of<br>distance of 2 meters<br>(53 dB) | .101 | 1 | .101 | 2.306 | .141 | .084 |
| N100 Latency of<br>distance of 2 meters<br>(53 dB) | 1.896 | 1 | 1.896 | .032 | .860 | .001 |

Univariate analyses were conducted to further explore the effects of group differences on the N100 component across varying auditory distances and intensities. The results revealed a marginal group effect on the N100 amplitude at a distance of 0.5 meters (65 dB), F(1, 25) = 3.965, *p* = .058, with a partial eta squared of .137, indicating a moderate effect size despite the non-significant p-value. This trend-level finding may suggest potential group-related differences in early auditory processing at closer spatial distances, warranting further investigation with larger samples. No significant group differences were observed for N100 latency at 0.5 meters (*p* = 1.000), nor for either amplitude or latency at 1 meter (59 dB) or 2 meters (53 dB), with all p-values well above the .05 threshold. The partial eta squared values for these comparisons ranged from .000 to .090, suggesting small effect sizes. Overall, the pattern of results implies that group-related modulation of early auditory ERP responses, particularly in amplitude, may be more pronounced in proximal acoustic condition

### N100 Responses to Direction

**Figure 6.**
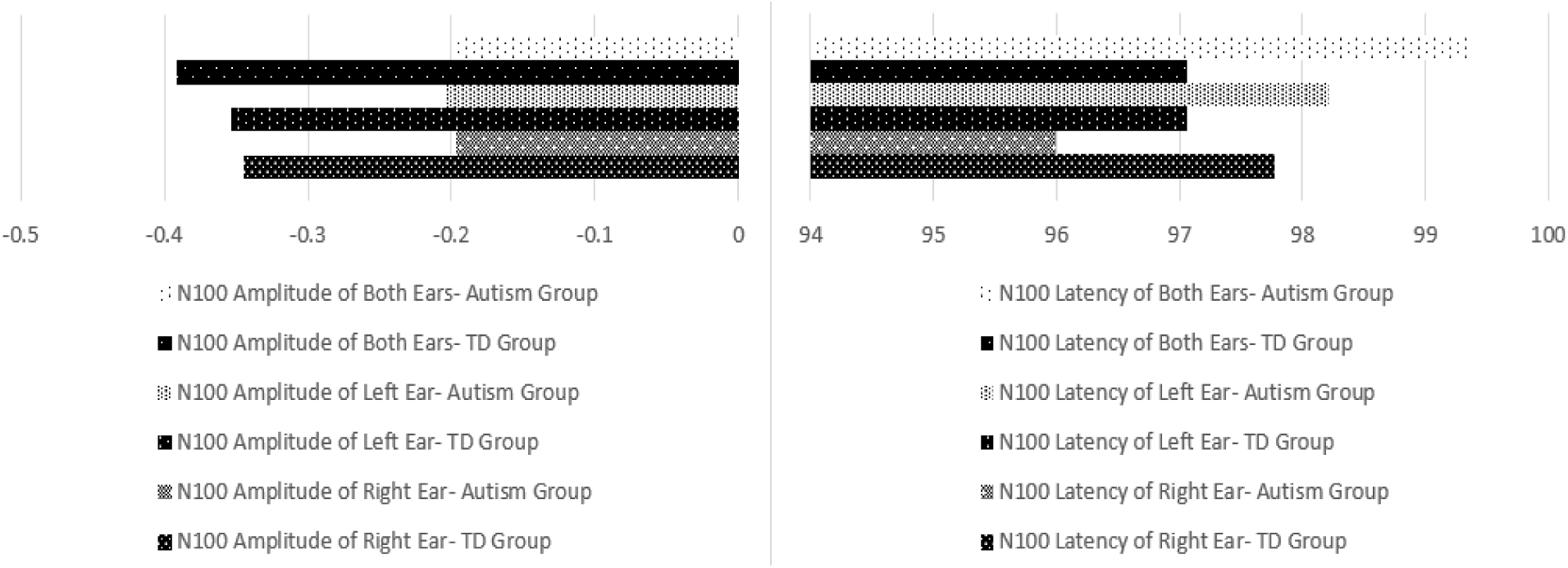
Comparison of N100 amplitude and latency between children with ASD and TD peers across three auditory spatial conditions (right, left, and bilateral).

Across all auditory spatial conditions (right, left, and binaural), the ASD group exhibited consistently reduced N100 amplitudes compared to TD peers (e.g., binaural: −0.39 µV TD vs. −0.20 µV ASD). In contrast, N100 latencies showed a more variable pattern: shorter latency for ASD in right ear stimulation (96 ms ASD vs. 97.8 ms TD) but longer latencies in left ear (98.2 ms ASD vs. 97.1 ms TD) and binaural conditions (99.3 ms ASD vs. 97.1 ms TD).

**Table 5.**
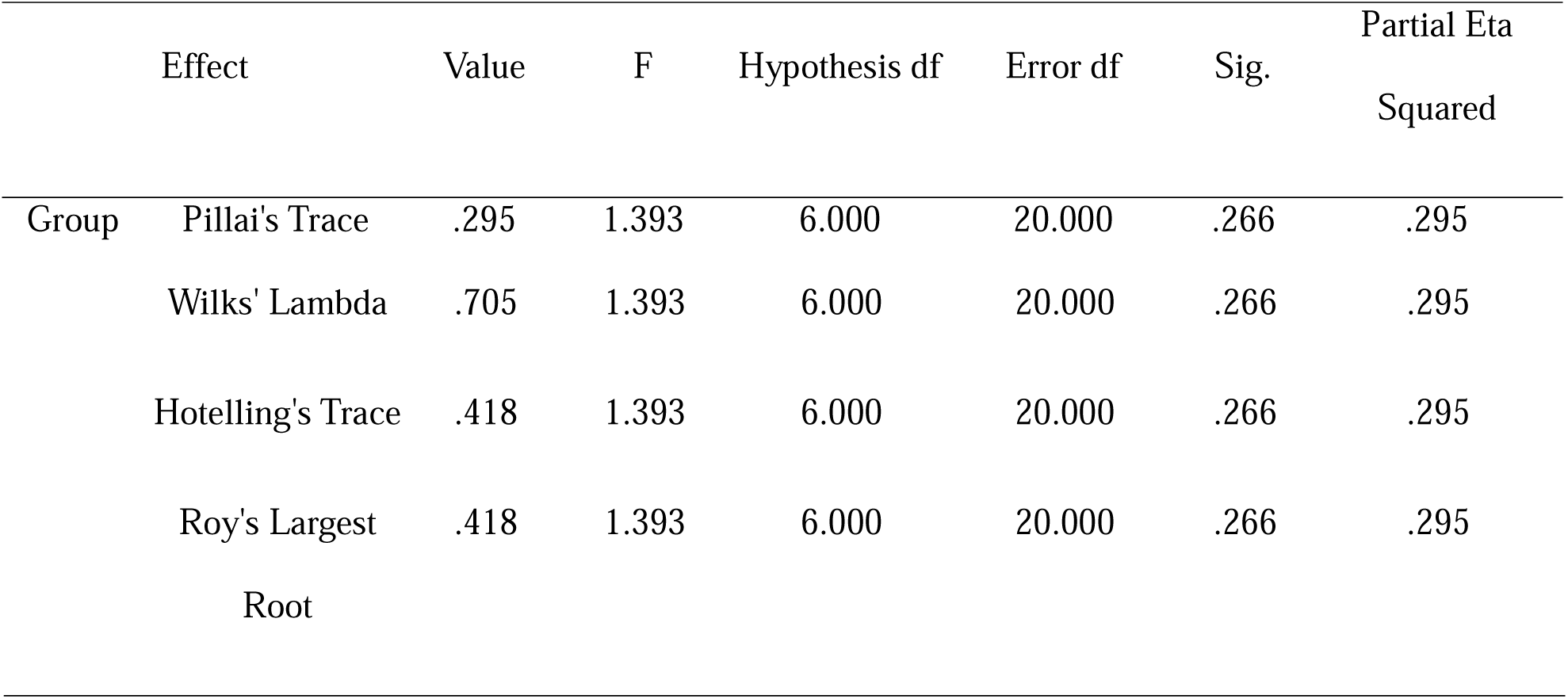
Summary of multivariate test results for the main effect of group on N100 amplitude and latency, considering all direction deviation conditions combined.

|  |  |  |  |  |  |  | Partial Eta |
| --- | --- | --- | --- | --- | --- | --- | --- |
|  | Effect | Value | F | Hypothesis df | Error df | Sig. | Squared |
| Group | Pillai's Trace | .295 | 1.393 | 6.000 | 20.000 | .266 | .295 |

|  |  |  |  |  |  |  |
| --- | --- | --- | --- | --- | --- | --- |
| Wilks' Lambda | .705 | 1.393 | 6.000 | 20.000 | .266 | .295 |
| Hotelling's Trace | .418 | 1.393 | 6.000 | 20.000 | .266 | .295 |
| Roy's Largest | .418 | 1.393 | 6.000 | 20.000 | .266 | .295 |
| Root |  |  |  |  |  |  |

A one-way multivariate analysis of variance (MANOVA) was conducted to examine group differences (ASD vs. Typically Developing) across six dependent variables. The multivariate effect of group was not statistically significant, Wilks’ Lambda = .705, F(6, 20) = 1.393, p = .266, partial η² = .295, indicating no strong overall group effect across the combined dependent variables.

However, the partial eta squared of .295 suggests a moderate effect size, which may indicate meaningful group-related variation, even in the absence of statistical significance. This trend-level multivariate effect warrants further investigation of univariate outcomes to explore possible specific group differences.

**Table 6.**
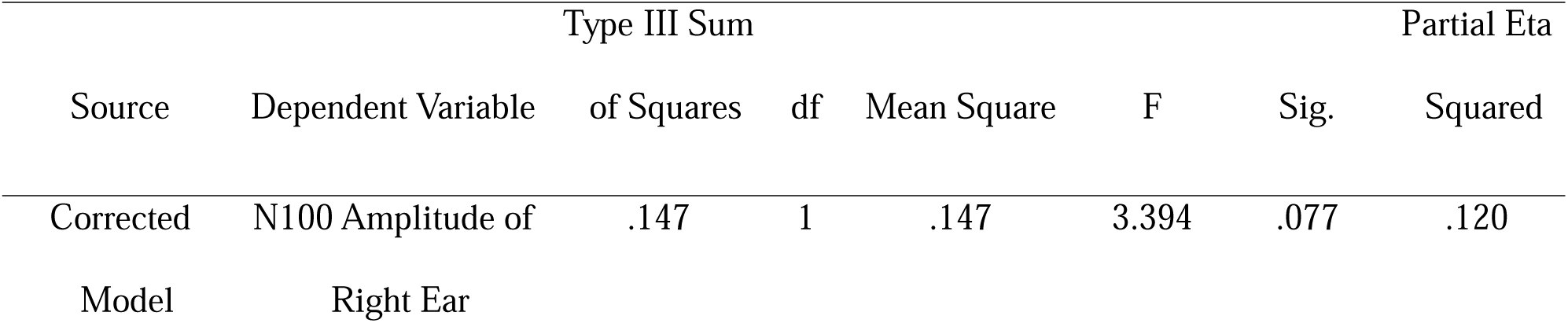
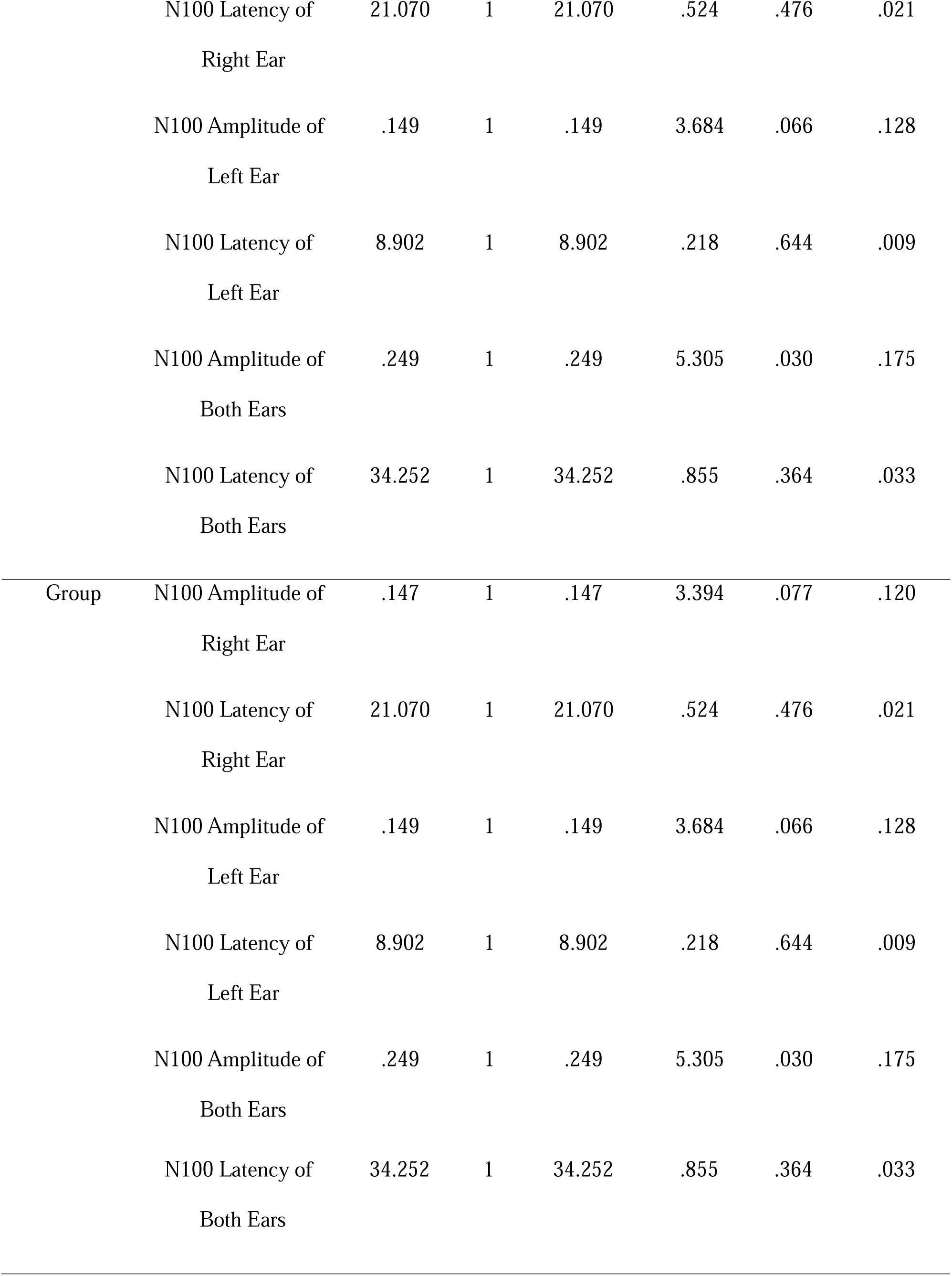
Tests of between-subjects effects on N100 amplitude and latency across directional conditions comparing children with ASD and TD peers.

| Source | Dependent Variable | Type III Sum |  |  | F | Sig. | Partial Eta Squared |
| --- | --- | --- | --- | --- | --- | --- | --- |
|  |  | of Squares | df | Mean Square |  |  |  |
| Corrected Model | N100 Amplitude of Right Ear | .147 | 1 | .147 | 3.394 | .077 | .120 |
|  | N100 Latency of | 21.070 | 1 | 21.070 | .524 | .476 | .021 |
|  | Right Ear |  |  |  |  |  |  |
|  | N100 Amplitude of | .149 | 1 | .149 | 3.684 | .066 | .128 |
|  | Left Ear |  |  |  |  |  |  |
|  | N100 Latency of | 8.902 | 1 | 8.902 | .218 | .644 | .009 |
|  | Left Ear |  |  |  |  |  |  |
|  | N100 Amplitude of | .249 | 1 | .249 | 5.305 | .030 | .175 |
|  | Both Ears |  |  |  |  |  |  |
|  | N100 Latency of | 34.252 | 1 | 34.252 | .855 | .364 | .033 |
|  | Both Ears |  |  |  |  |  |  |
| Group | N100 Amplitude of | .147 | 1 | .147 | 3.394 | .077 | .120 |
|  | Right Ear |  |  |  |  |  |  |
|  | N100 Latency of | 21.070 | 1 | 21.070 | .524 | .476 | .021 |
|  | Right Ear |  |  |  |  |  |  |
|  | N100 Amplitude of | .149 | 1 | .149 | 3.684 | .066 | .128 |
|  | Left Ear |  |  |  |  |  |  |
|  | N100 Latency of | 8.902 | 1 | 8.902 | .218 | .644 | .009 |
|  | Left Ear |  |  |  |  |  |  |
|  | N100 Amplitude of | .249 | 1 | .249 | 5.305 | .030 | .175 |
|  | Both Ears |  |  |  |  |  |  |

|  |  |  |  |  |  |  |
| --- | --- | --- | --- | --- | --- | --- |
| N100 Latency of | 34.252 | 1 | 34.252 | .855 | .364 | .033 |
| Both Ears |  |  |  |  |  |  |

A series of univariate ANOVAs was conducted to examine the effect of group (Autism vs. Typically Developing) on N100 amplitude and latency in response to auditory stimuli presented to the right ear, left ear, and binaurally. The analysis revealed a statistically significant group effect for N100 amplitude in the binaural condition, F(1, 25) = 5.305, p = .030, with a partial eta squared of .175, indicating a moderate effect size. This finding suggests that children with autism exhibit significantly reduced N100 amplitudes compared to their typically developing peers when auditory input is presented to both ears simultaneously, highlighting potential impairments in early bilateral auditory integration.

In contrast, while the group effects for N100 amplitude in the right ear (F = 3.394, p = .077, η² = .120) and left ear (F = 3.684, p = .066, η² = .128) did not reach statistical significance, both approached the conventional threshold (p < .05) and showed moderate effect sizes. These near-significant trends may indicate underlying group differences in lateralized auditory processing that merit further exploration in studies with increased statistical power.

With regard to N100 latency, none of the group comparisons reached significance across right (p = .476), left (p = .644), or binaural (p = .364) conditions. Partial eta squared values for latency ranged from .009 to .033, reflecting small effect sizes. These results suggest that, while the timing of early auditory responses may not differ significantly between groups, the strength (amplitude) of cortical responses—especially during binaural processing, is more sensitive to group-related differences.

## Discussion

This study provides novel neurophysiological evidence highlighting atypical early-stage auditory processing in children with autism indexed by the N100 event-related potential (ERP) component. Utilizing ecologically valid speech stimuli, we systematically manipulated auditory dimensions of pitch, intensity (operationalized via source distance), and spatial direction. Across all auditory conditions, children with ASD exhibited consistently reduced N100 amplitudes relative to TD peers, while N100 latency remained largely preserved. This pattern suggests that although the timing of early auditory cortical responses is intact, the neural signal strength is attenuated, supporting predictive coding frameworks and models of reduced sensory gain in autism (Lawson et al., 2014; Lieder et al., 2019; Palmer et al., 2017).

### N100 and Pitch

The most robust group difference emerged in response to normal pitch stimuli, with children with ASD demonstrating significantly diminished N100 amplitudes compared to TD controls (M = −0.198 µV vs. −0.392 µV; F(1,25) = 5.305, p = .030, η² = .175). This medium-to-large effect aligns with prior electrophysiological research indicating pitch perception deficits in ASD (Russo et al., 2008; Zhang et al., 2019; Ong et al., 2024) and may reflect underlying cortical hypo-responsiveness or atypical synaptic integration (Baranek et al., 2013; Simon et al., 2017; Carroll et al., 2021). Notably, the normal pitch condition—characterized by low perceptual salience— elicited the weakest neural response in the ASD group, emphasizing reduced sensitivity to baseline auditory stimuli (Edgar et al., 2015).

In response to low-pitch stimuli, a trend-level reduction in N100 amplitude was observed (F = 2.992, p = .096, η² = .107), suggesting possible frequency-specific encoding anomalies. Although not statistically significant, the moderate effect size and consistent directionality of the effect (−0.21 µV vs. −0.35 µV) may indicate atypical tonotopic organization (Kujala et al., 2013; Bharadwaj et al., 2022) or diminished top-down attentional modulation (Orekhova et al., 2012). Given the importance of low-pitched cues in prosody and affective speech, these sensory processing deficits could underlie the social-communicative challenges frequently reported in autism.

### N100 and Intensity (Sound Distance)

Analysis of N100 responses across varying intensities and source distances revealed a consistent pattern of reduced amplitudes in the ASD group, especially to proximal stimuli (0.5 m, 65 dB), with near-significant differences (F(1,25) = 3.97, p = .058, η² = .137). Although multivariate analyses did not yield significant group differences overall (Pillai’s Trace = 0.198, p = .565), univariate trends indicate early cortical hypoactivation in ASD during processing of physically salient sounds. These findings resonate with predictive coding models positing attenuated precision weighting of sensory inputs and weakened prediction error signals in ASD (Crasta et al., 2021; Lieder et al., 2019).

From an ecological perspective, reduced neural responsiveness to proximal, intense stimuli may impede the ability to orient toward relevant social and environmental cues—capacities crucial for joint attention, social referencing, and language acquisition (Sharda et al., 2015; Wood et al., 2019). The absence of latency differences suggests that temporal processing remains intact, highlighting the specific impairment in response magnitude rather than timing.

### N100 and Direction

Binaural stimulation elicited the strongest group differences, with children with ASD showing significantly reduced N100 amplitudes compared to TD peers (F(1,25) = 5.305, p = .030, η² = .175). This reduction likely reflects impaired binaural integration, possibly localized to brainstem structures such as the superior olivary complex (Kulesza Jr. et al., 2011; ElMoazen et al., 2020). These neural deficits may contribute to difficulties in spatial speech perception, speaker localization, and segregation of speech from background noise—common challenges for autistic children in complex auditory environments (da Silva Mayerle et al., 2023; Soskey et al., 2017).

Additionally, unilateral conditions (right ear: F = 3.394, p = .077, η² = .120; left ear: F = 3.684, p = .066, η² = .128) showed trend-level reductions in N100 amplitude in ASD, indicating possible disruptions in hemispheric lateralization or selective auditory encoding (Herringshaw et al., 2016; Li et al., 2023). These near-significant findings underscore the importance of investigating asymmetrical cortical processing in ASD further. Consistent with earlier results, latency remained unaffected across all directional conditions, reinforcing that neural timing is preserved despite amplitude attenuation (Groen et al., 2009; Zhou et al., 2022).

### Clinical and Theoretical Implications

The observed amplitude reductions in the N100 component during passive listening tasks, even without explicit behavioral demands, highlight the automaticity of early sensory impairments in ASD. Given its reliability and ease of elicitation in passive paradigms, including in minimally verbal or young children, the N100 holds promise as a noninvasive biomarker for atypical auditory processing. Portable, cost-effective EEG systems like OpenBCI could facilitate scalable screening tools in community and school settings, potentially identifying children at risk for language and social communication difficulties before observable behavioral manifestations.

From a developmental psychopathology standpoint, these results align with accumulating evidence that disruptions in early sensory encoding may precede and contribute to downstream challenges in language, social reciprocity, and cognition (Baranek et al., 2013; Schultz-Krohn, 2021). However, this study’s focus on high-functioning boys necessitates replication in more diverse cohorts, including females and individuals with varying cognitive abilities. Longitudinal designs will be critical to ascertain whether early N100 anomalies predict later developmental outcomes.

## Conclusion

In conclusion, this study demonstrates that early auditory cortical processing, as indexed by the N100 amplitude, is consistently attenuated in children with ASD across multiple auditory domains, including pitch, intensity, and spatial direction. These findings reveal foundational sensory encoding disruptions that may cascade into higher-order communicative and social impairments characteristic of autism. The preservation of N100 latency alongside amplitude attenuation highlights a specific deficit in neural gain or sensory precision rather than temporal processing speed.

Moreover, the consistent amplitude reductions across ecologically valid auditory conditions underscore the potential utility of the N100 as a scalable neural biomarker for early identification and intervention in ASD. Future research expanding beyond male, high-functioning populations, incorporating additional ERP components, behavioral correlates, and longitudinal follow-ups, will be essential to fully elucidate the developmental and clinical significance of these neural differences.

### Limitations and Future Directions

This study included exclusively male participants to reduce variability in verbal ability, as females, both with and without ASD, typically demonstrate stronger language skills and different processing profiles. Therefore, caution is advised when generalizing these findings to autistic girls, whose auditory processing patterns may differ. Furthermore, despite sample size limitations, participants in the two groups were matched for age, intelligence, native language, socioeconomic status, culture, handedness, and level of functioning (ASD level 1), and had no history of comorbid neurological or psychiatric conditions. The history of receiving therapeutic interventions (such as speech therapy and occupational therapy) was also taken into account to minimize the influence of environmental and educational factors and facilitate interpretation of neurological differences. This was done to maximize internal validity. However, larger and more diverse samples and longitudinal designs would further enhance generalizability and clinical relevance.

## Key Points

- Children with Autism Spectrum Disorder (ASD) show reduced N100 amplitudes in response to speech sounds, reflecting atypical early-stage auditory processing.
- This study reveals consistent—but often subtle—amplitude reductions across pitch, intensity (distance), and spatial direction, with trend-level effects suggesting pervasive neural differences.
- Temporal aspects of auditory processing (N100 latency) remain intact in ASD, indicating specific deficits in sensory gain rather than timing.
- These findings support predictive coding theories proposing reduced sensory precision in ASD, particularly during passive listening.
- Portable EEG combined with ecologically valid speech stimuli holds promise for developing noninvasive biomarkers for early identification and intervention in children with ASD.

## Acknowledgments

The authors gratefully acknowledge the valuable support and assistance of Dr. Morteza Izadifar, Dr. Diana Sînziana Duca, Cristina Lemeni, Tiberiu Ciortan, Roxana Toderean and the Star of Hope Autism Center in this research.

## Statements and Declarations

### Funding

This research received no external funding.

### Conflict of Interest

None of the authors has any potential conflicts of interest to disclose.

### Ethics approval

All procedures involving human participants were conducted in accordance with the ethical standards of the institutional research committee and the 1964 Helsinki Declaration and its later amendments. The study protocol was reviewed and approved as exempt by the Ethics Committee of the University of Tabriz (IR.TABRIZU.REC.1403.172).

### Consent to publish

Written informed consent for publication was obtained from all participants or, where applicable, from their parents or legal guardians.

### Consent to Participate

Written informed consent was obtained from all individual participants. For minors, consent was obtained from their parents or legal guardians.

### Availability of Data, Materials, and/or Code

The datasets and code used in the current study are available from the corresponding author on reasonable request.

### Authors’ Contributions

**Sara Sharghilavan**: Conceptualization, Methodology, Investigation, Formal Analysis, and Writing – Original Draft.

**Leila Mehdizadeh Fanid**: Supervision, Conceptualization, Methodology, Writing – Review & Editing.

**Oana Geman**: Project Administration, Investigation, Resources, Software, and Formal Analysis.

**Hassan Shahrokhi**: Methodology and Validation.

**Hadi Seyedarabi**: Software and Formal Analysis.

## Notes

### Competing Interest Statement

The authors have declared no competing interest.

